# Pulsatile contractions and pattern formation in excitable actomyosin cortex

**DOI:** 10.1101/2021.02.22.432369

**Authors:** Michael F. Staddon, Edwin M. Munro, Shiladitya Banerjee

## Abstract

The actin cortex is an active adaptive material, embedded with complex regulatory networks that can sense, generate, and transmit mechanical forces. The cortex exhibits a wide range of dynamic behaviours, from generating pulsatory contractions and travelling waves to forming organised structures. Despite the progress in characterising the biochemical and mechanical components of the actin cortex, the emergent dynamics of this mechanochemical system is poorly understood. Here we develop a reaction-diffusion model for the RhoA signalling network, the upstream regulator for actomyosin assembly and contractility, coupled to an active actomyosin gel, to investigate how the interplay between chemical signalling and mechanical forces regulate stresses and patterns in the cortex. We demonstrate that mechanochemical feedback in the cortex acts to destabilise homogeneous states and robustly generate pulsatile contractions. By tuning active stress in the system, we show that the cortex can generate propagating contraction pulses, form network structures, or exhibit topological turbulence.

## Introduction

The actin cortex is an adaptive active material that dynamically regulates its mechanical properties to maintain or change cell shapes [1–4]. Actin cortex can display a wide range of dynamic behaviours from driving intracellular pulsatory contractions [5, 6] to cellular-scale polarized flows [5, 7, 8] and assembling protrusive or contractile structures during cell motility and cytokinesis [9, 10]. These behaviours must adapt to the cell’s local environment and developmental stage. For instance, cellular-scale pulsatile contractions are often observed during developmental morphogenesis, where pulsatile contractions act as a mechanical ratchet to sequentially alter cell size and shapes [11–14], leading to tissue bending or elongation [15]. At the intracellular level, actomyosin pulses occur with chaotic spatiotemporal dynamics [5, 16, 17]. Travelling waves of actomyosin contraction can propagate across the cortical surface [16, 18], and can be highly organised as in the case of surface contraction waves, which propagate as single waves from one pole of the cell to the other [19, 20]. In other physiological contexts, stable contractile structures are needed, as in the formation of stress fibers during cell adhesion [10], assembly of actomyosin purse-string during wound healing or gap closure [21–23], or the formation of a contractile ring during cytokinesis [24, 25]. While the biochemical pathways underlying actomyosin dynamics are known, the mechanisms by which actomyosin-driven mechanical forces feedback to upstream chemical signals to govern patterns and flows in the actin cortex remain poorly understood [4].

Alan Turing, in his seminal paper, demonstrated that reaction-diffusion systems can autonomously generate a wide variety of spatiotemporal patterns observed in nature, but noted that mechanical forces may also play an important role in pattern formation [26]. In recent years, purely biochemical models have been proposed for actomyosin pattern formation. RhoA and its downstream effectors of actin and myosin have been shown to form an activator-inhibitor system that exhibits excitable dynamics and oscillations [17, 18, 27]. Here autocatalytic production of the activator RhoA leads to delayed accumulation of the inhibitor F-actin, producing local pulses of RhoA activity. Since F-actin diffusion is negligible compared to RhoA, this system produces travelling pulses of RhoA activity but cannot generate Turing patterns [18, 27]. By tuning RhoA production rates locally, static patterns of RhoA can be generated [27]. This raises the question if actomyosin patterns in the cortex strictly rely on biochemical cues or can spontaneously and robustly emerge via interplay between mechanical forces and biochemical signaling.

Mechanochemical feedback in the cytoskeleton is another mechanism for generating spatiotemporal patterns [28–31]. Active gel models of the cytoskeleton have suggested that pulsatory patterns can emerge from contractile instabilities driven by a positive feedback, in which active stress drives advective flows of stress producing factors such as actin and myosin [30, 32, 33]. These contractile instabilities cluster myosin into a few high concentration regions, but regulating myosin contraction with an independent RhoA oscillator can prevent the collapse of myosin and sustain oscillations [34]. However recent studies suggest a negative feedback loop between actomyosin and RhoA [17, 18, 27], raising the question of how the feedback between RhoA and actomyosin mechanics are regulated to generate flows and patterns. Additionally, it remains unclear how the same molecular system can regulate the formation of stable contractile structures [4, 35] or exhibit turbulent dynamics [16].

Here we develop a reaction-diffusion model for an excitable signalling network comprising RhoA and actomyosin, embedded in an active mechanical medium, to study how the feedback between biochemical signalling and mechanical stresses regulate mechanochemical patterns and flows in the actin cortex. We specifically ask how pulsatory flows arise from the coupling between a fast diffusing activator (RhoA) and a slow diffusing inhibitor (actomyosin), how mechanochemical feedbacks stabilise contractile instabilities into localised patterns, and how active stresses propagate local contractile signals or drive turbulent dynamics. Our model builds upon recent experimental observations of activator-inhibitor relationship between RhoA and actomyosin [17], by introducing myosin-driven contraction to provide a mechanical feedback. By tuning just two biologically relevant parameters, the basal rate of RhoA production and the magnitude of active contractile stress, our model is able to capture a wide range of dynamic phases observed in the actomyosin networks, from travelling waves and pulsatile contractions to stable contractile networks and turbulent dynamics. We find that mechanochemical feedback acts to destabilise stationary states to robustly generate pulsed contractions. At high contractile activity and low basal rates of RhoA production, stable patterns of actomyosin emerge. Furthermore, the mechanochemical system encodes memory of transient perturbations that allows local signals to be translated into propagating contraction waves or stable patterns.

## Results

### Mechanochemical feedback generates robust pulsatile contractions

To elucidate the role of mechanochemical feedback in the generation of dynamic behaviours and instabilities in the actin cortex, we first study an ordinary differential equation model for the coupling between RhoA, actomyosin, and mechanical strain (Fig. 1a). Here for simplicity, we neglect spatial variations in chemical concentrations and study the coupled dynamics of RhoA and actomyosin in a locally homogeneous region of the cortex. Recent experiments on *Xenopus* oocytes [18] and *C. elegans* embryos [17] suggest that actomyosin pulsation in the cortex is regulated by the excitable dynamics of RhoA GTPase – the upstream regulator of actomyosin assembly and force production. Local autocatalytic activation of RhoA drives rapid initiation of RhoA pulses, followed by F-actin assembly and myosin recruitment. As actomyosin concentrations increase, F-actin dependent accumulation of the RhoA GTPase-activating proteins RGA-3/4 terminate the pulse through a delayed negative feedback [17].

**Figure 1.**
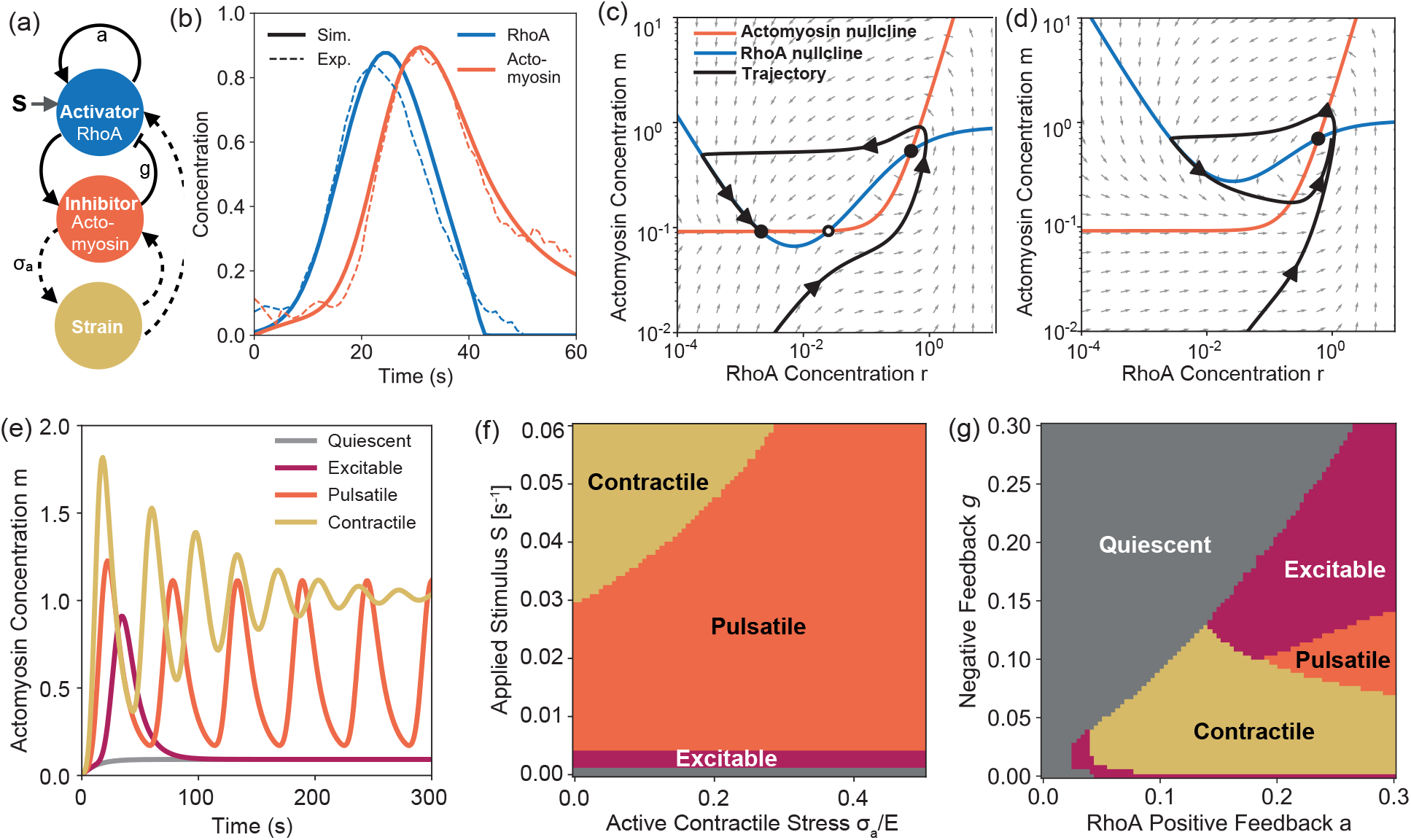
Mechanical feedback sustains pulsatile contractions in excitable active medium. (a) Feedback loop schematic of the system. Solid lines indicate biochemical feedback. Dashed lines indicate mechanical feedback. (b) RhoA (blue) and actomyosin concentration over time in the excitable phase during a single pulse. Dashed lines indicated experimentally measured values, solid lines show model with best fit parameters. (c-d) Trajectory curves and nullclines showing the fixed points for RhoA (blue) and actomyosin (orange) in the excitable (b) and in the pulsatile regime (c). Black arrows on the trajectory are equally spaced in time. Grey arrows show motion in phase space. Solid circles represent stable fixed points and the open circle is the unstable fixed point. (e) Concentrationof actomyosin over time in the quiescent phase (*S* = 0), excitable phase (*S* = 0.002 s^−1^), pulsatile phase (*S* = 0.025 s^−1^), and contractile phase (*S* = 0.075 s^−1^). (f) Phase diagram of the system, for varying contractile stress, *σ*_*a*_*/E*, and applied stimulus, *S*, with *η/E* = 5s. (g) Phase diagram of the system, for varying rate of autocatalytic production of RhoA, *a*, negative feedback parameter, *g*. See Table I in Supplemental Material for a list of default parameters in the model.

Here, for simplicity, we represent F-actin and myosin as a single species - actomyosin - whose properties combine active force production (by myosin) and inhibition of RhoA (by F-actin). This representation is justified since myosin-II mostly co-localizes with F-actin in the cortex. Based on previous experimental observations [17], we make the following assumptions in our model. First, RhoA is activated at a constant basal rate, *S*. Second, active RhoA promotes further production of active RhoA via an autocatalytic feedback, and also promotes the production of actomyosin. Third, actomyosin (F-actin) promotes local inactivation of RhoA by recruiting GAPs. With these assumptions, the rate of production of RhoA-GTP can be written in terms of the RhoA-GTP concentration *r* and actomyosin concentration *m*:

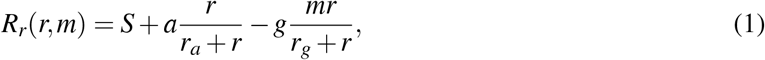

where *S* is the applied stimulus (or basal rate of RhoA production) representing the activity of Rho-GEF, *a* is the rate of autocatalytic production of RhoA, and *g* is a negative feedback parameter arising from F-actin-driven accumulation of the GAP RGA-3/4 that inactivates RhoA. The constants *r*_*a*_ and *r*_*g*_ represent the threshold concentrations of RhoA above which the rate of RhoA production is independent of RhoA concentration. The rate of actomyosin production is given by:

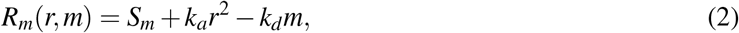

where *S*_*m*_ is the basal rate of actomyosin production, *k*_*a*_ is the actomyosin assembly rate and *k*_*d*_ is the disassembly rate. While the model parameters *S, S*_*m*_, *k*_*a*_ and *k*_*d*_ are directly avalable from single-molecule data [17], the rest are calibrated by fitting our model to the experimental data for the concentration of RhoA and actomyosin during one contraction pulse (Fig. 1b).

To introduce mechanical feedback into this model, we describe a locally homogeneous region of the cortex as an active viscoelastic material with strain *u*, with contractile stresses generated by the actomyosin:

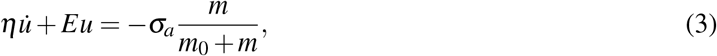

where *E* is the compressional elastic modulus, *η* is the viscosity, *σ*_*a*_ (> 0) is the maximum active stress arising from actomyosin-driven contractions, and *m*_0_ is the concentration of actomyosin at half-maximum stress. The viscoelastic time-scale, *τ* = *η/E* and relative contractility *σ*_*a*_*/E* are estimated from available data on actin cortex of *C. elegans* [36].

Mechanical stress feeds back to the dynamics of both RhoA and actomyosin through conservation of mass; if the size of the system doubles then the concentration of chemical species must halve. Written explicitly, *∂*_*u*_(*c*(1 + *u*)) = 0, which implies that *∂*_*u*_*c* = −*c/*(1 + *u*), where *c* is the chemical concentration. Thus, we have the following governing equationss for the coupled dynamics of RhoA and actomyosin:

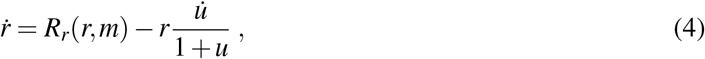

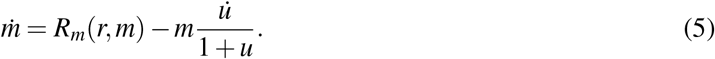

In the above equations, actomyosin provides additional feedback to both RhoA and itself through changes in mechanical strain. An increase in local actomyosin concentration induces a local contraction, which in turn increases the actomyosin concentrations. By contrast, when actomyosin concentration decreases, there is local strain relaxation leading to a decrease in actomyosin concentration. Using the parameters calibrated from experiments, we simulated these dynamics numerically [37], for different values of the activity parameters *S* and *σ*_*a*_, to understand the role of mechanochemical feedback in actin cortex dynamics.

The model yields four distinct dynamic behaviours – quiescent, excitable, pulsatile, and contractile, depending on the magnitude of the applied stimulus, *S* (Fig. S 1). For low *S*, the system displays an excitable behaviour – a single pulse of RhoA, followed by a pulse of actomyosin and contractile strain buildup, before reaching a steady-state (Fig. 1e). This excitable behaviour can be visualised in phase space as a trajectory about the intersecting nullclines (Fig. 1c). For this excitable pulse, we observed a loop about the high (*r,m*) fixed point before settling to a steady state at the lower fixed point.

As the applied stimulus is increased, other behaviours emerge (Fig. 1e-f). For a small increase in *S*, we observe sustained pulsatile contractions, when the RhoA nullcline shifts up (Fig. 1d), resulting in a single fixed point at high concentrations and the system is trapped in a limit cycle. At even higher values of *S*, the limit cycle is unstable and the pulse amplitude decays until the system settles in a contracted state with high actomyosin concentration and strain. Finally, for very low applied stimulus, we observe a quiescent mode; both RhoA and actomyosin steadily increase to a fixed set-point.

While pulsatile contractions are observed in the absence of contractile stress (*σ*_*a*_ = 0), we find that including mechanical feedback helps to sustain oscillations over a wider range of the parameter space (Fig. 1f). As actomyosin concentration drops after a pulse, the mechanical strain relaxes, further reducing both actomyosin and RhoA concentrations away from a fixed point, allowing another pulse to occur. These pulsatile states occur above a threshold value for the autocatalytic positive feedback parameter *a* and for moderate values of the negative feedback parameter *g* (Fig. 1g). When the negative feedback is higher than the positive feedback we observe quiescent behaviour, where any changes in RhoA concentration are quickly slowed down by the negative feedback and brought to its equilibrium value. When *a* is high compared to *g* we observed several distinct regions of dynamic behaviours. When both *a* and *g* are low, the system settles to a contractile steady-state. When both *a* and *g* are high the system is excitable, and when *a* is high and *g* is moderate the system becomes pulsatile (Fig. 1g).

### Spatial propagation of pulsatile flows and contractile pattern formation

To investigate the role of active mechanical stresses and RhoA signalling on spatial patterns and flows within the actin cortex, we develop a continuum active gel model of the actin cortex, coupled to the excitable RhoA signalling network (Fig. 1a). Using the *C. elegans* embryo as a model system, we develop a one-dimensional description of the actin cortex with periodic boundary conditions, assuming azimuthal symmetry around the long axis of the cell. The actin cortex is modelled as a Maxwell viscoelastic material [36], behaving elastically at short times and remodelling over longer time scales, with flows in the cortex advecting RhoA and actomyosin. The constitutive equation defining the time-evolution of stress field *σ* (*x,t*) is given by:

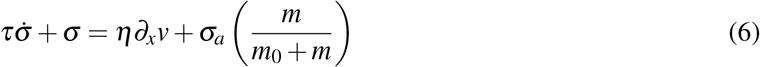

where the spatial coordinate *x* denotes distance along the surface of the cell, *τ* is the viscoelastic relaxation time scale, *η* is the viscosity, and *v*(*x,t*) is the cytoskeletal flow velocity. The second term on the right hand side of the above equation is the active stress term with *σ*_*a*_ the active contractile stress, and *m*_0_ is the actomyosin concentration at half-maximum active stress. Local balance of viscoelastic forces with cytosolic drag and actomyosin-generated active contractile forces can be written as:

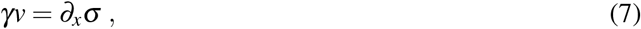

where *γ* is the frictional drag coefficient. By taking a spatial derivative in Eq. (6) and using Eq. (7) we obtain the following equation for the velocity field:

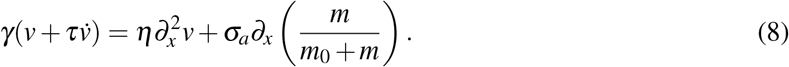

Mechanical feedback is then introduced through myosin-induced F-actin flows which advect both RhoA and myosin, leading to their local accumulation due to convergent flows or depletion via divergent flows. These advective flows compete with reaction and diffusion of RhoA and actomyosin:

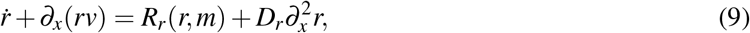

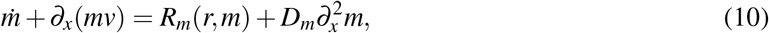

where *D*_*r*_ and *D*_*m*_ are the diffusion coefficients for RhoA and actomyosin, taken from Nishikawa et al [34]. The default parameter for active stress and viscoelastic time scale are taken from Saha et al [36]. The model equations are then numerically integrated [38] in a periodic box of length *L* = 10*λ*, where 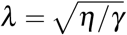 is the hydrodynamic length scale.

As shown in Fig. 2a-c, the numerical solutions predict a wide diversity of dynamic states as the active contractility 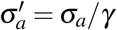 and RhoA stimulus *S* are varied– from stationary patterns to propagating waves and pulsatile flows (Fig. S 2). At low 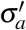, diffusion dominates over advection, giving rise to spatially uniform concentration profiles that exhibit excitable, oscillatory, or contractile dynamics (Fig. 2c, top row), as observed in the cellular-scale model (Fig. 1). In the absence of mechanical feedback, reaction-diffusion alone cannot generate spatial patterns because the activator, RhoA, diffuses much faster than the inhibitor, actomyosin (*D*_*r*_ ≫ *D*_*m*_). In this regime, the homogeneous state becomes unstable and Turing patterns emerge, only when *D*_*r*_ ≪ *D*_*m*_ (Fig. S 3).

**Figure 2.**
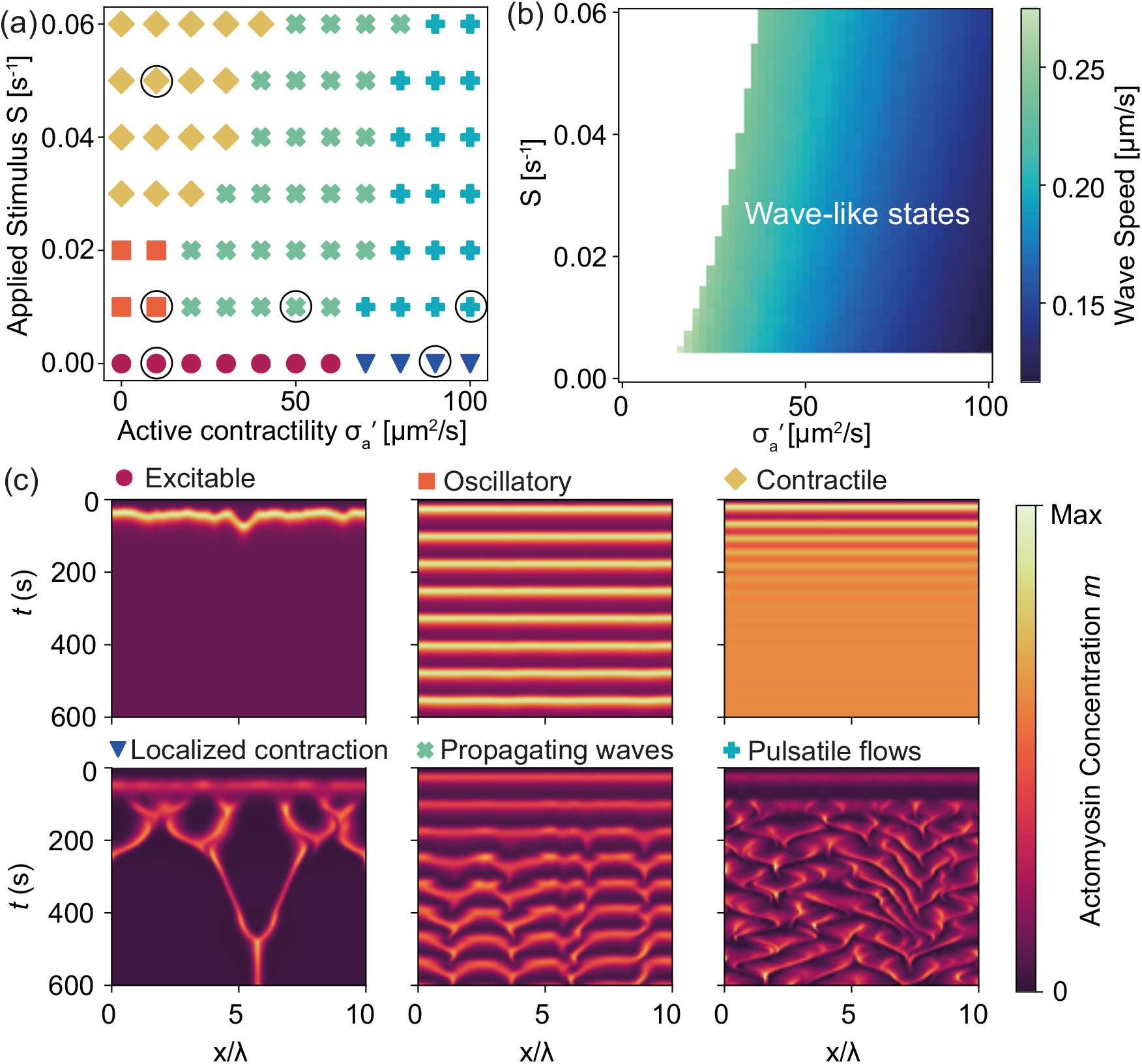
Active stress generates spatial patterns and pulsatory flows. (a) Phase diagram of the dynamic states of the system as a function of applied Rho stimulus *S* and active contractility 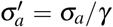. Encircled data points correspond to kymographs in panel (c). (b) Actomyosin wave speed, computed from linear stability analysis of the model equations, for varying *S* and 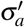. (c) Kymograph of actomyosin concentration in different regimes (left to right, top to bottom): excitable, oscillatory, homogeneous contractile, localised contractions, propagating waves and pulsatile flows. See Supplemental Tables I and II for a list of model parameters.

As 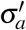 is increased, contractile instabilities develop due to local accumulation of actomyosin, allowing finite wavelength patterns to emerge. At low *S* and high 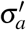, we observe stable localized peaks of actomyosin and RhoA (Fig. 2c, bottom left). Initially, autocatalytic positive feedback of RhoA creates small RhoA concentration peaks (Fig. 3a, 1st panel), which in turn produce actomyosin (Fig. 3a, 2nd panel). As actomyosin begins to accumulate, large inward flows are generated, further increasing both RhoA and actomyosin concentrations (Fig. 3a, 3rd panel). Finally, actomyosin-induced inhibition of RhoA results in RhoA localization on either side of the actomyosin peak (Fig. 3a, 4th panel), in contrast to Turing patterns where activators and inhibitors overlap. With no RhoA advection, RhoA diffuses away from the peak, producing actomyosin behind it and generates a travelling wave (Fig. S 4a). With no actomyosin advection, actomyosin concentrations remain too low to prevent RhoA from diffusing and create a uniform steady state (Fig. S 4b). When several contractile actomyosin foci exist, they attract each other and merge into a single peak. This phase is reminiscent of equatorial RhoA zones during cell division [39, 40], and medial [41] and junctional [42] RhoA domains in polarized epithelia.

**Figure 3.**
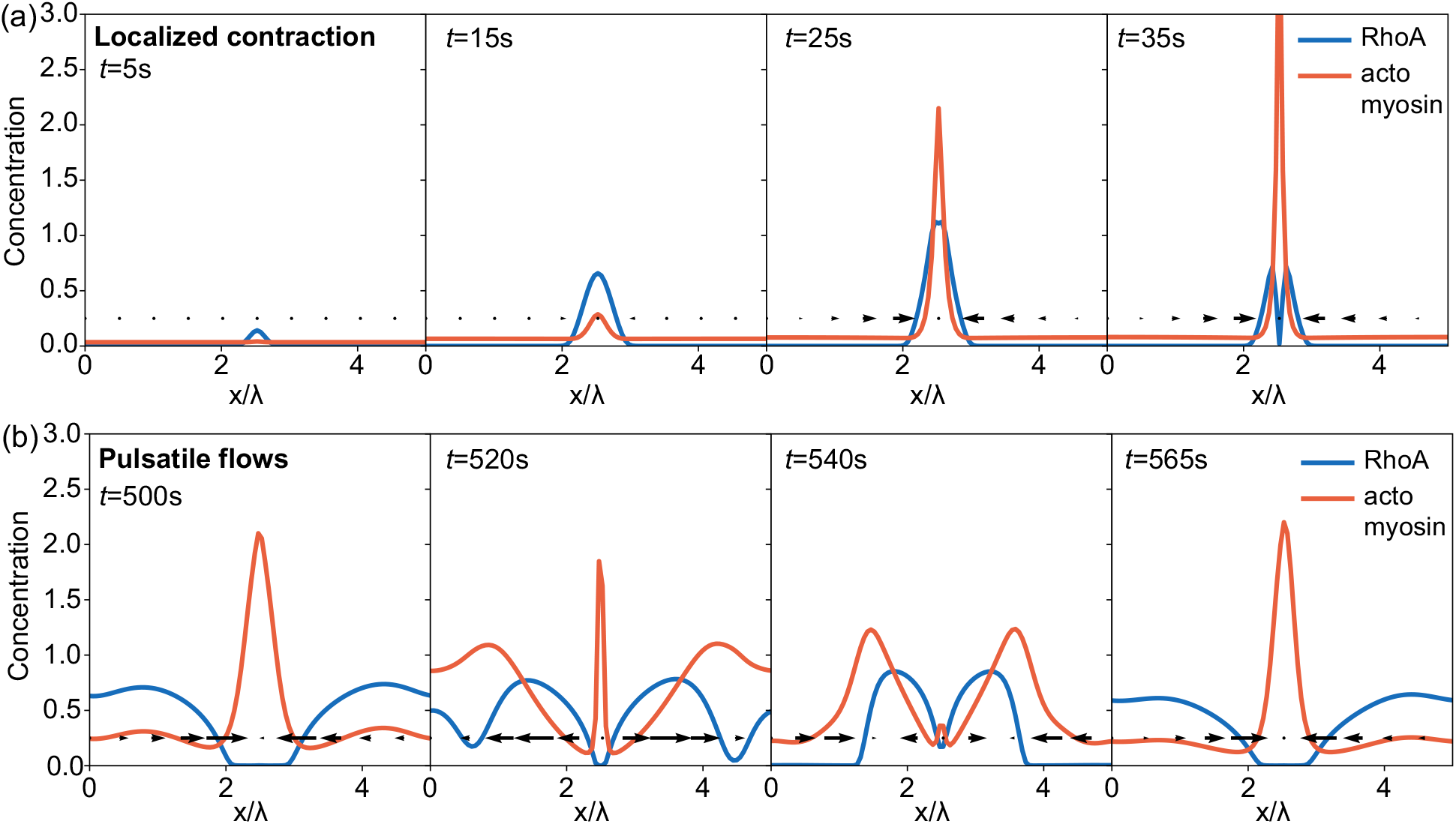
Feedback mechanisms for pulsatile flows and stationary pattern formation. (a) RhoA and actomyosin concentration during localized contraction 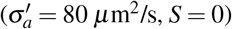 at t = 5s, 15s, 25s, and 35s (left to right). (b) RhoA and actomyosin concentration profiles over a pulsatile flow cycle 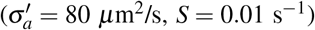 at t = 500s, 520s, 540s, and 565s (left to right). Arrows indicate flow velocity.

At higher *S* and moderate 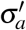, we observe propagating waves and pulsatile flows (Fig. 2c, bottom). A higher level of excitation in RhoA generates a localized actomyosin peak with high contractility. Away from the actomyosin peak, RhoA concentrations are higher and are advected towards actomyosin (Fig. 3b, 1st panel). Advected RhoA produces actomyosin as it moves, such that the newly assembled actomyosin generates flows away from the centre, reducing the actomyosin concentration at the centre (Fig. 3a, 2nd panel). Once the initial contraction dissipates, the two remaining actomyosin peaks merge, completing a cycle (Fig. 3a, panels 3-4).

In the propagating waves states, we observe the periodic formation of RhoA pulses, which travel in waves before annihilating as two waves meet, as observed in the starfish oocyte [16, 18]. As 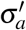 is increased, actomyosin pulses generate large contractile flows that advect neighbouring pulses, creating chaotic, aperiodic motion. The pulses are highly localised, with small regions of high actomyosin concentration that form before dispersing, followed by a new pulse elsewhere, as in the *C. elegans* embryo [17]. Mechanical feedback through advection is necessary for the waves to form (Fig. S 4d). Without advection of RhoA however, we may still observe waves, since the actomyosin generated by RhoA forms clusters that travel with RhoA (Fig. S 4c). However, these waves display much less chaotic motion when compared to the system with advection (Fig. 2c, bottom right).

Linear stability analysis of the continuum model reveals the role of active stress in destabilising the homogeneous state of the system (Fig. 2b, Fig. S 5). At low *S*, three fixed points exist (Fig. 1b), with the lower fixed point being stable for high 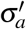. As *S* is increased, the RhoA nullcline shifts up until only one fixed point remains and the system enters the pulsatile regime (Fig. 1c). Increasing 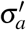 leads to contractile instabilities that manifest as propagating waves and pulsatile flows (Fig. 2c). While active stress is required for wave propagation, higher 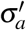 leads to lower wave speeds (Fig. 2b).

### Response to local bursts of contractile activity

While our model can capture many of the dynamic states observed in the actin cortex, cells must have the ability to actively switch between flowing and contractile states during physiological transitions. Such state transitions may be triggered by a local up-regulation in RhoA activity, which may be induced in response to mechanical forces or by cell cycle checkpoints. But how mechanochemical coupling shapes the spatiotemporal response to transient localized changes in RhoA activity remains unclear.

To understand how RhoA signals can propagate through space and induce state transitions, we locally turned on RhoA activity and examined the output response. Starting from rest with no applied stimulus (*S* = 0), we applied an increased stimulus at the central region (*S* = 0.03 s^−1^) (Fig. 4a). By changing active stress in the system, we were able to regulate both the ability for the input RhoA to propagate in space, and for the system to remember the spatial location of the signal. We observed three distinct phases (Fig. 4a): (i) propagation of a bistable front, where the memory of the signal location is lost and a global increase in RhoA concentration is observed, (ii) a soliton phase with transient spatial memory, and (iii) a high memory phase where a stable RhoA pattern is maintained.

**Figure 4.**
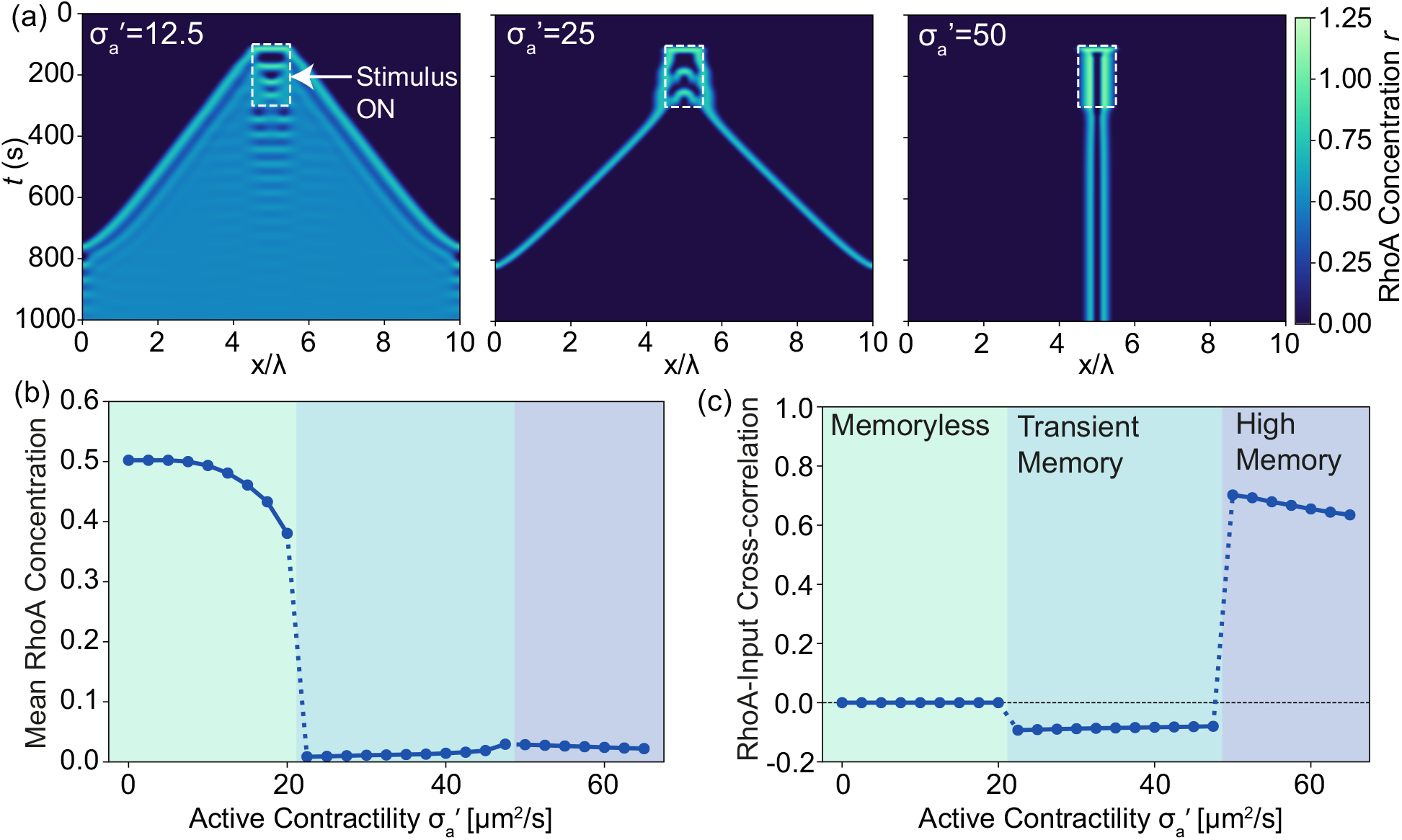
Response to local bursts of contractile activity. (a) Kymograph of Rho concentration upon local transient application of Rho stimulus: *S* = 0.03 s^−1^ inside box (dashed rectangle) and *S* = 0 outside box, for different values of active stress: (left) bistable front propagation for 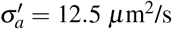, (middle) solitons for 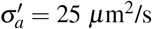, and (right) localised contraction for 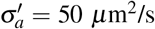. (b) Spatially averaged RhoA concentration, 300s after the application of stimulus. (c) Correlation between RhoA concentration and input stimulus *S*(*x, t*), averaged over the last 120s.

At low active stress 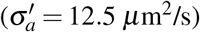, RhoA is excited into a pulsatile state within the activation region, while spreading laterally through diffusion (Fig. 4a, left). A front of highly concentrated RhoA travels away from the source, increasing the total RhoA and actomyosin in the system (Fig. 4b). This is reminiscent of a bistable front propagation in classical excitable systems [42, 43], where the system switches to the high concentration stationary state (Fig. 2c).

As active stress is increased, we find that mechanical feedback is able to tune the properties of the biochemical system away from classical excitable systems. At moderate active stress 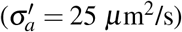, RhoA pulses spread out as two solitons before annihilating as they meet (Fig. 4a, middle), as seen in surface contraction waves [19, 20]. In contrast to classical excitable systems, actomyosin generated behind the RhoA wavefront increases its own concentration through contractile flows. This region of highly concentrated actomyosin behind the wave acts as a barrier that inhibits RhoA, preventing the system from switching to the high fixed point, and instead creates a soliton. Such a barrier can be seen at 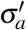, although it is too weak to prevent diffusion, with the band of reduced RhoA concentration behind the wavefront (Fig. 4a, left).

At high active stress 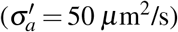, the contractile forces within the activated region are strong enough to maintain a spatially localized state that persists after the activation, remaining stationary in a fixed location at the centre of activation (Fig. 4a, right). This suggests a potential mechanism for cells to direct the locations of contractility. Without the guidance of externally induced RhoA activation, the system in this parameter regime is incapable of spontaneously forming a stable pattern.

To quantify the input-output relationship of the system, we measured the correlation between the input signal and the output RhoA concentration at long times (Fig. 4c). At low 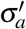, RhoA spreads outwards, leading to a loss of correlation between input and the output signal, akin to a memoryless system. For the soliton case, the shape of the input signal is remembered, resulting in a negative correlation as the waves travel away from the source. Finally, in the spatially patterned phase, a high-memory state emerges where RhoA remains localized at the centre of activation, with a strong positive correlation between the input and the output. These results suggest that mechanical stresses can play an important role in biochemical signal propagation, and in retaining the spatial memory of activity. For low active stress, signals propagate the fastest with global changes in contractility and no memory of the spatial location of the signal. As active stress is increased, we observe contraction waves propagating away from the source, displaying transient memory. At higher stresses, a high memory state develops, where transient local RhoA activations create localized contractile states.

### Network formation, pulsatile flows and topological turbulence

We now proceed to analyze our model in two spatial dimensions to investigate how contractile patterns form and propagate on the surface of a cell. The equation governing the temporal evolution of the velocity field **v** is given by

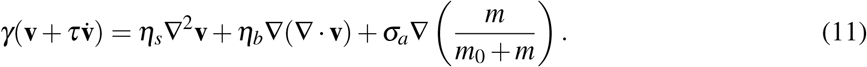

where **v** is the velocity vector, *η*_*s*_ and *η*_*b*_ are the shear and bulk viscosities. We assume that the bulk viscosity is higher than the shear viscosity, such that 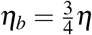 and 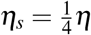, where *η* is the experimentally measured viscosity [34]. The reaction-diffusion equations for RhoA and actomyosin concentration fields are given by

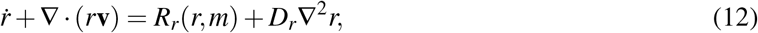

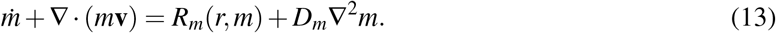

In the two-dimensional model, as we vary active stress (*σ*_*a*_) and the applied stimulus of RhoA (*S*) we observe similar dynamic phases as in the one-dimensional model, though with more varied spatial organisation. At low applied stimulus and high active stress, we observe the formation of stable contractile networks (Fig. 5a, Movie 1), analogous to the pattern formation phase observed in the one-dimensional model. Here, an initial excitable pulse forms many small cluster of highly concentrated actomyosin (Fig. 5a, left panel). Over time, the clusters stretch and merge with other nearby clusters (Fig. 5a, middle panel), eventually creating a space-spanning network of actomyosin, which gradually condenses into a stable configuration of fully connected edges and vertices (Fig. 5a, right panel). These condensates of actomyosin are stabilized by contractile flows, similar to the localized contraction peaks observed in the one-dimensional model (Fig. 3a). This sequence of actomyosin patterning is stringly reminiscent of apical microridge formation in some epithelial cells [44].

**Figure 5.**
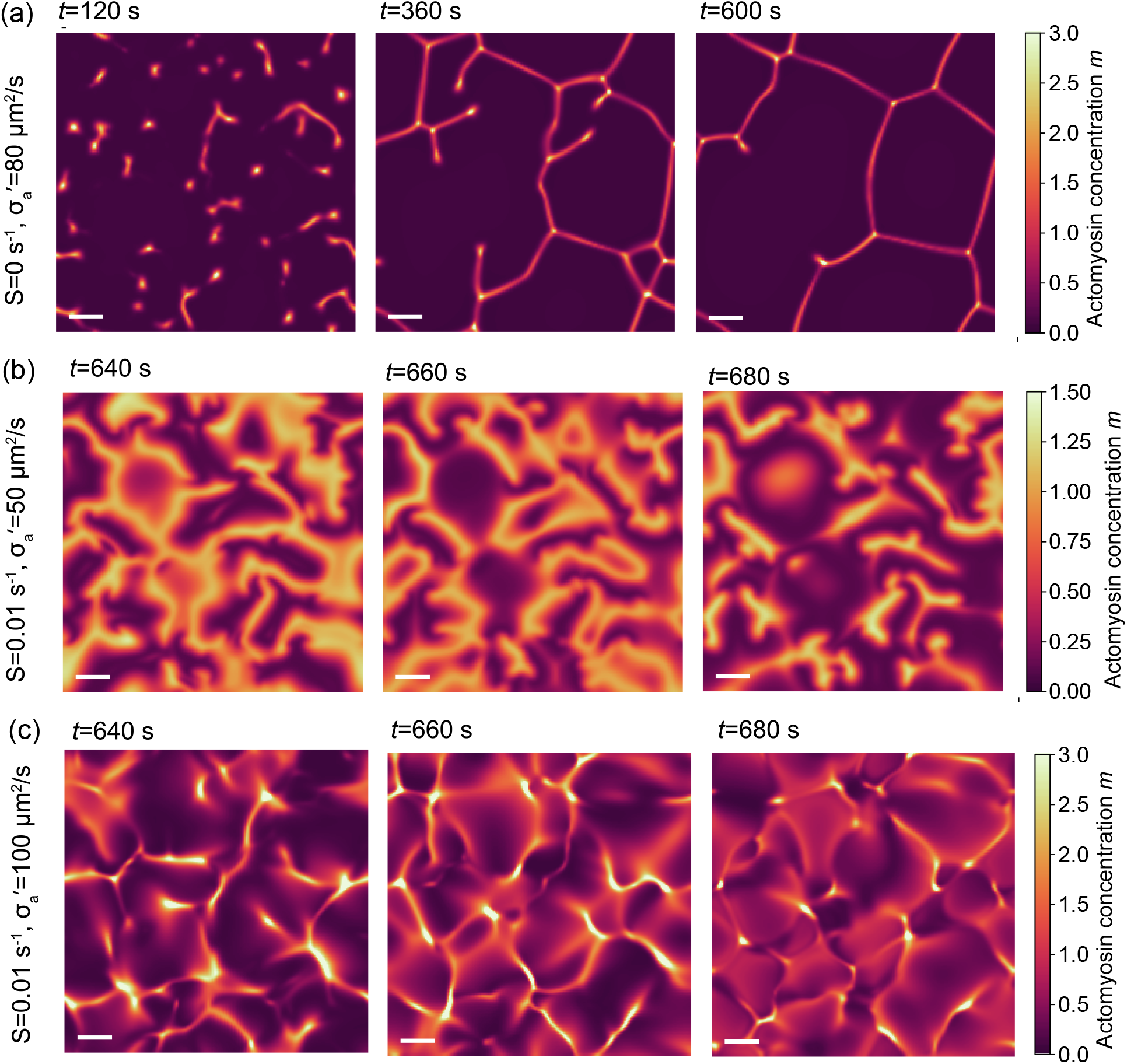
Pattern formation and pulsatile flows in two spatial dimensions. (a-c) Actomyosin concentration field obtained from simulations of the active gel model in two spatial dimensions, showing (a) contractile network formation 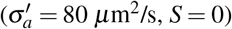, (b) propagating waves 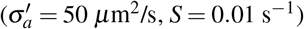 and (c) localized pulsatile contractions 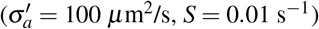. Scale bar indicates a distance of *λ* = 15 *µ*m.

As the applied stimulus *S* is increased we observe pulsatile contractions and propagating waves of RhoA and actomyosin. At moderate active stress we observe propagating waves (Fig. 5b, Movie 2), in which waves of actomyosin propagate with constant velocity until two waves collide and annihilate, with new waves periodically produced. Finally, at high applied stimulus and high active stress, we observe the pulsatile contraction phase (Fig. 5c, Movie 3). Pulses of actomyosin contract into highly concentrated foci before dispersing and new pulses are formed. These dynamics are distinct from propagating waves as the pulses of actomyosin move more erratically as clusters merge, and show much more spatial variation in concentration.

The propagating waves of RhoA show features of topological turbulence, as recently reported in the starfish oocyte [16]. In this regime, RhoA and actomyosin exist with a limit cycle (Fig. 1d), allowing us to extract the phase of oscillation using the relation 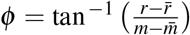, where 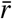 and 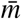 are the mean RhoA and actomyosin concentrations (Fig. 6a). We find that numerous topological defects exist in the phase field, indicating locations where the phase field is discontinuous. These may be parameterised by the winding number: the total amount that the phase changes by following an anti-clockwise contour around the defect. From the phase velocity field **v**_*ϕ*_= ∇*ϕ*(Fig. 6b), the integer winding number is calculated as 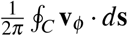, where *C* is a closed curve around the defect containing no others. In a +1 defect, the phase field spirals anti-clockwise about the defect, while in a -1 defect it spirals in a clockwise direction (Fig. 6d). In our model, we find approximately the same number of +1 and -1 defects. In addition, oppositely charged defects attract one another, and annihilate when close enough, conserving the overall charge of the system (Movie 4).

**Figure 6.**
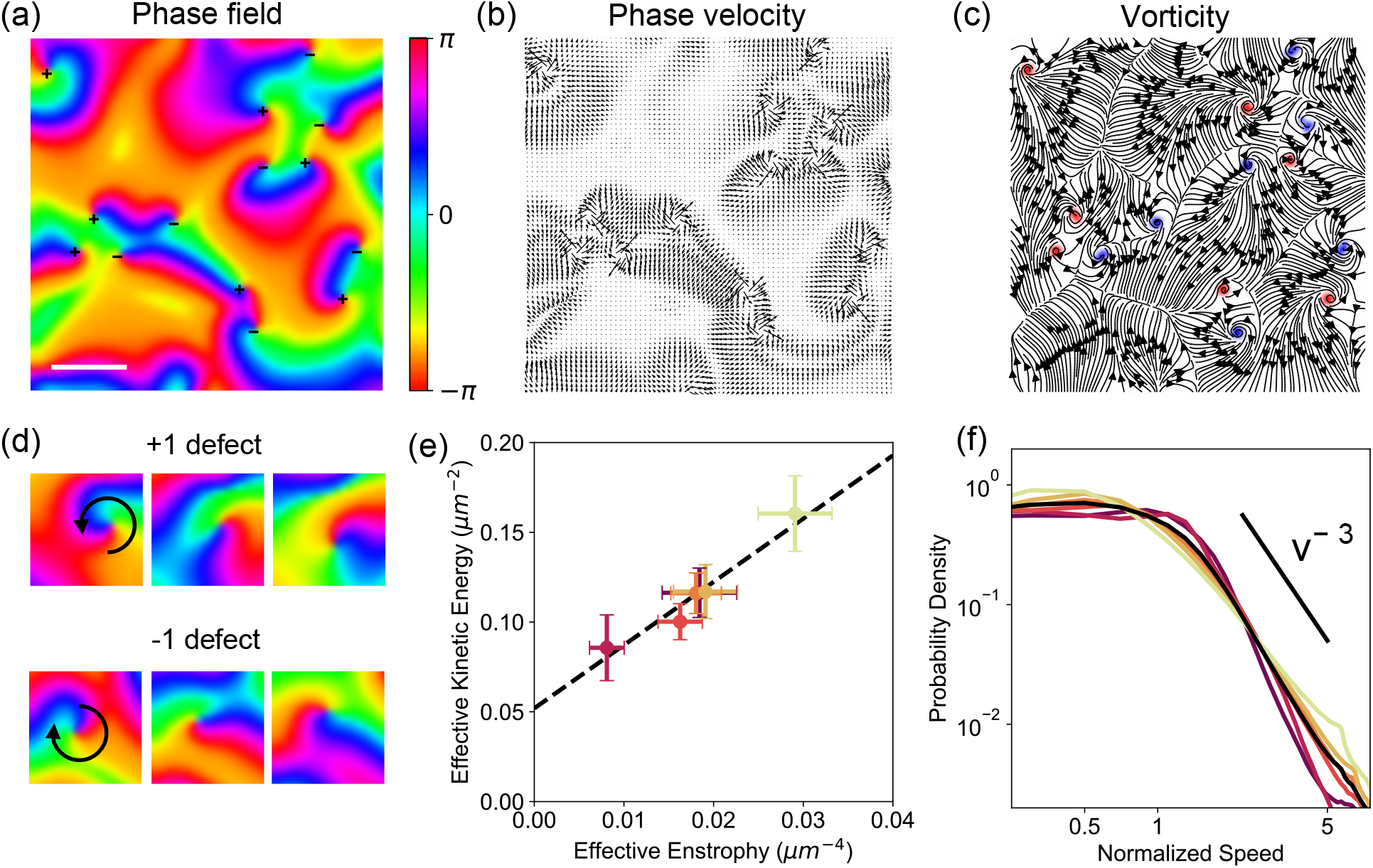
Topological turbulence in RhoA flows. (c) The RhoA phase field, *ϕ*, corresponding to the dynamics in Fig. 5b. Plus and minus symbols represent the location of +1 and -1 defects, respectively. (b) Spatial map of the phase velocity field **v**_*ϕ*_= ∇*ϕ*. (c) Spatial map of contours of the phase vorticity field *ω* = ∇ × **v**_*ϕ*_. Red indicates positive vorticity, corresponding to plus defects. Blue indicates negative vorticity, corresponding to minus defects. Lines indicate streamlines of the phase velocity field. (d) Representative images of a +1 defect (top) and -1 defect (bottom) over an oscillation cycle at (left to right) *t* = 640s, 660s, and 680s. (e) Linear scaling between the effective kinetic energy and the effective enstrophy of the RhoA phase velocity. (f) Probability density of normalized speed of RhoA phase field motion, for different values of active stress, 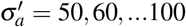. Light colours indicate higher magnitudes of active stress.

The phase velocity also describes an effective velocity field due to a potential flow with point vortices of magnitude *ω* = ∇ *×* **v**_*ϕ*_, with +1 and -1 defects corresponding to the center of vorticies with clockwise or anticlockwise rotations. We find a linear scaling between the effective kinetic energy 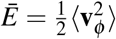 and the effective enstrophy 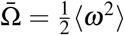, a measure of vorticity, for different values of active stress, in agreement with experimental data on RhoA in starfish oocytes [16]. Furthermore, when we quantify the phase velocity statistics, we find a power law tail in the probability density function which decays like *v*^−3^, likely due to interacting vortices, consistent with experimental results [16] and theoretical predictions for vortex interactions [45]. Thus our model can quantitatively capture the key statistics and scaling laws of defect mediated turbulence in the RhoA phase field.

## Discussion

Chemical signalling and mechanical forces are both essential to regulate the dynamics of the cellular actin cortex. Here we demonstrate that mechanochemical coupling of actomyosin and its upstream regulator RhoA are necessary to generate waves, patterns and pulsatile dynamics in the actin cytoskeleton. Mechanical feedback helps to robustly sustain pulsatile contractions, and provides a new mechanism for spatial patterning of RhoA and myosin, distinct from classical Turing patterns [26]. Having an excitable cortex with tunable dynamics and mechanochemical feedback can be beneficial in a number of morphogenetic contexts. Sustained pulsed contractions can act as a mechanical ratchet to sequentially reduce cell area and drive tissue bending [12–14, 46–48]. Mechanochemical feedback may be important in enabling robust control of this morphogenetic ratchet. Cells may also display excitable behaviours which are governed by mechanics, for example, with active contractions triggered by large mechanical strains on the cell, helping to prevent tissue rupture [49, 50].

At low RhoA activity and high contractile activity, myosin forms stationary patterns, with RhoA localised on either side of the myosin peak. In two spatial dimensions, we observe the formation of many small clusters of high myosin concentrations, which stretch towards neighbouring clusters and condense into a stable contractile domains of actomyosin and RhoA. This is reminiscent of stable patterns observed *in vivo* on the apical surfaces of epithelial cells, including microridges [44], stable junctional and medial RhoA domains [41, 42]. As contractile activity is increased, the waves of myosin become highly peaked, creating pulsing contractile networks, reminiscent of pulsatile contractions in the *C. elegans* cortex [5]. At moderate activity, we observe periodic waves of myosin which annihilate before new wavefronts are generated in regions with low actomyosin concentration. These waves show features of topological turbulence, reminiscent of topological turbulence in RhoA concentration waves observed in starfish eggs [16]. Besides generating patterns and flows, the level of active contractile stress regulates the system’s response to local RhoA signals, enabling phase transitions and memory entrainment as the activity is increased. Together, these results highlight the importance of considering both mechanical forces and chemical reactions when modelling the actin cortex, and they reveal a variety of ways in which cells can tune the dynamic coupling between RhoA activity, force production, and advective transport to control morphogenetic behaviours.

## Author Contributions

SB, MFS and EM designed and developed the model. MFS performed model simulations and analytical calculations. SB, MFS and EM wrote the paper.

## Acknowledgements

SB acknowledges funding from the Royal Society (URF/R1/180187), Human Frontiers Science Program (RGY0073/2018) and the National Institutes of Health (R35GM143042). EM acknowledges funding from the National Institutes of Health (R01GM098441)

