## Supporting Information for "Pulsatile contractions and pattern formation in excitable actomyosin cortex"


*Linear Stability Analysis*.– To understand how mechanical stress and biochemical stimulus work together to generate pulsatile behaviour and propagating waves, we perform linear stability analysis about the homogenous steady-state (or fixed point) to determine the stability of the state, and the onset of oscillatory instabilities. In particular, we use linear stability analysis to compute the frequency, wave length and wave speed of the wave-like states that emerge from perturbations about quiescent homogeneous state. A perturbation of the form $\widetilde{y}e^{ikx}$ about the steady state, $(r, m, 0)$, results in the following Jacobian:

$$\widetilde{J} = \begin{pmatrix} \frac{aA}{(A+r)^2} - \frac{gmG}{(G+r)^2} - D_r k^2 & -\frac{gr}{(G+r)} & -ikr \\ 2k_a r & -k_d - D_m k^2 & -ikm \\ 0 & ik\frac{\sigma_a m_0}{\gamma\tau(m_0+m)^2} & -\frac{\gamma+\nu k^2}{\gamma\tau} \end{pmatrix}. \quad (1)$$

By finding the eigenvalue with the highest real part over all $k$, $\lambda(k^*)$, we obtain the fastest growing mode with wavelength $1/2\pi k^*$, frequency $\text{Im}(\lambda(k))$, and a wave speed of $\text{Im}(\lambda(k))/2\pi k^*$.

TABLE I. Default Biochemical parameters

| Parameter | Value |
|---|---|
| $S$ | 0 $\mathrm{s}^{-1}$ |
| $a$ | 0.1609 $\mathrm{s}^{-1}$ |
| $n$ | 1 |
| $r_a$ | 0.3833 |
| $g$ | 0.1787 $\mathrm{s}^{-1}$ |
| $r_g$ | 0.01 |
| $S_m$ | 0.0076 $\mathrm{s}^{-1}$ |
| $k_a$ | 0.1408 $\mathrm{s}^{-1}$ |
| $k_d$ | 0.0828 $\mathrm{s}^{-1}$ |

TABLE II. Default Mechanical parameters

| Parameter | Value |
|---|---|
| $\tau$ | 5s |
| $\lambda$ | 14.3 $\mu$m |
| $\sigma_a/\gamma$ | 49.8 $\mu\mathrm{m}^2\mathrm{s}^{-1}$ |
| $D_r$ | 0.1 $\mu\mathrm{m}^2\mathrm{s}^{-1}$ |
| $D_m$ | 0.01 $\mu\mathrm{m}^2\mathrm{s}^{-1}$ |

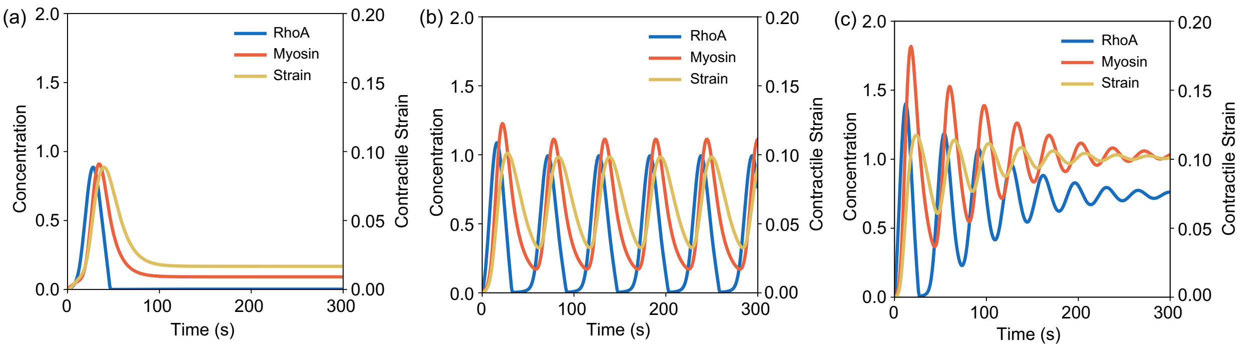


**Figure S** 1. **Dynamics in the activator-inhibitor mechanochemical model in response to changes in RhoA stimulus S.** (a-c) Dynamics of RhoA concentration (blue), actomyosin concentration (red), and contractile strain (yellow) in the (a) excitable phase ($\sigma_a/E = 0.2$, $S = 0.002$ $\mathrm{s}^{-1}$), (b) pulsatile phase ($\sigma_a/E = 0.2$, $S = 0.025$ $\mathrm{s}^{-1}$), and (c) the contractile phase ($\sigma_a/E = 0.2$, $S = 0.075$ $\mathrm{s}^{-1}$).

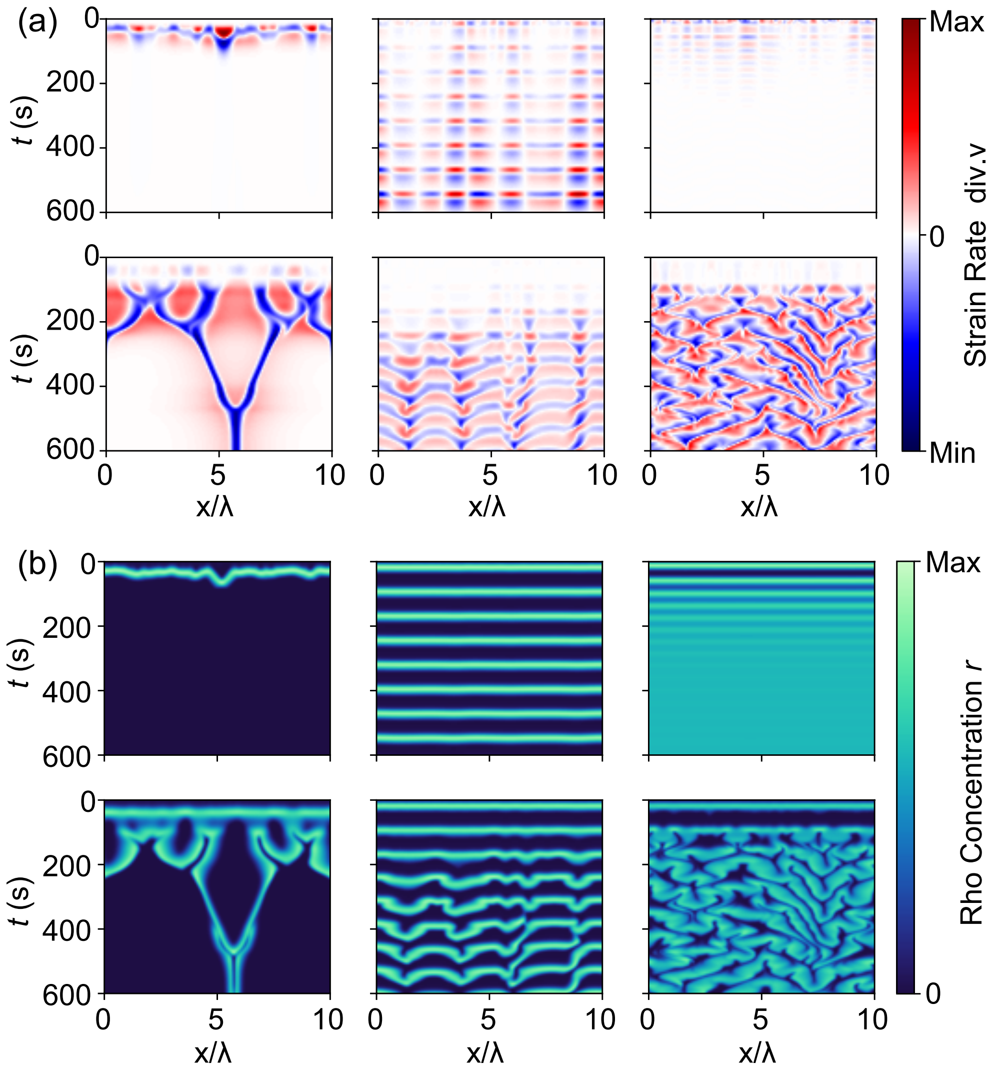


**Figure S** 2. **Strain rate and RhoA kymographs in different phases of the continuum active gel model.** Kymographs of (a) strain rate ($\partial_x v$), and (b) RhoA concentration for the different phases in the active gel model corresponding to the actomyosin kymographs shown in Fig. **??**c

.

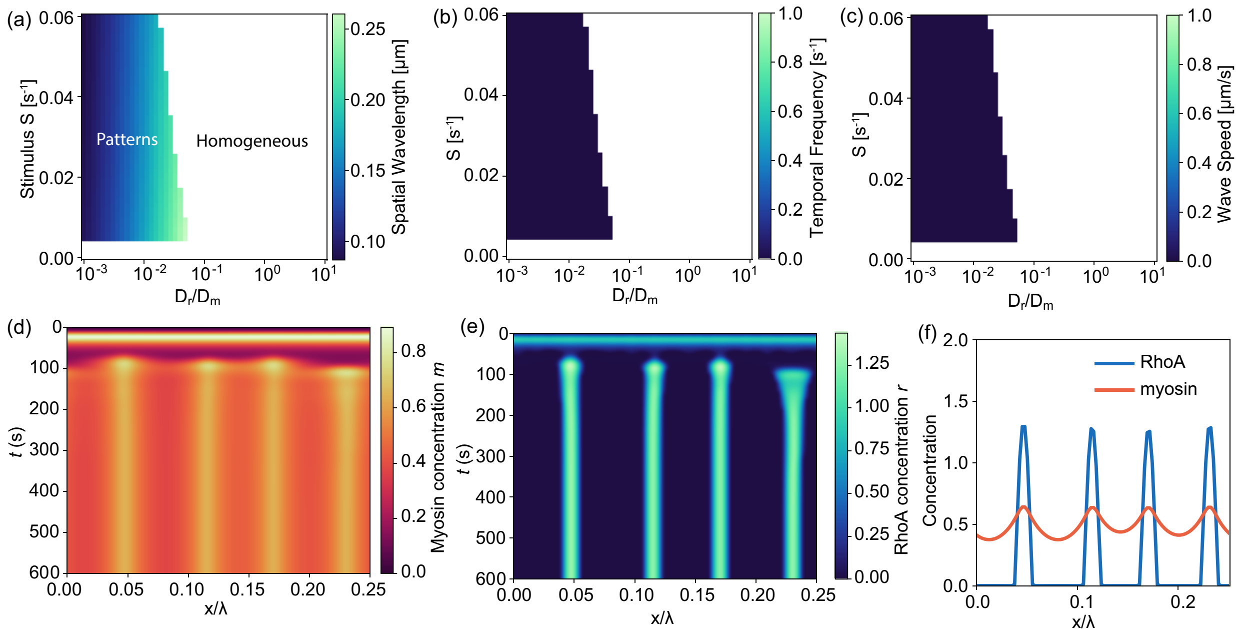


**Figure S 3. Reaction-diffusion alone is not sufficient to generate pulsatile flows and waves in RhoA-myosin systems.** (a) Spatial wavelength, (b) temporal frequency, and (c) wave speed computed by linear stability analysis of the RhoA-myosin reaction-diffusion model with $\sigma_a' = 0$, for varying stimulus $S$ and activator to inhibitor diffusivity ratio, $D_r/D_m$. White regions in the parameter space show homogeneous steady states. Default parameter value of $D_r/D_m$ is 10. (d-e) Kymographs of actomyosin concentration (d) and RhoA concentration (e) showing pattern formation for low RhoA diffusivity ($D_r/D_m = 0.01$, $S = 0.01$ s$^{-1}$). (f) Steady-state spatial profiles of RhoA and actomyosin concentrations for the patterns corresponding to (d) and (e). Overlapping peaks indicates Turing pattern formation.

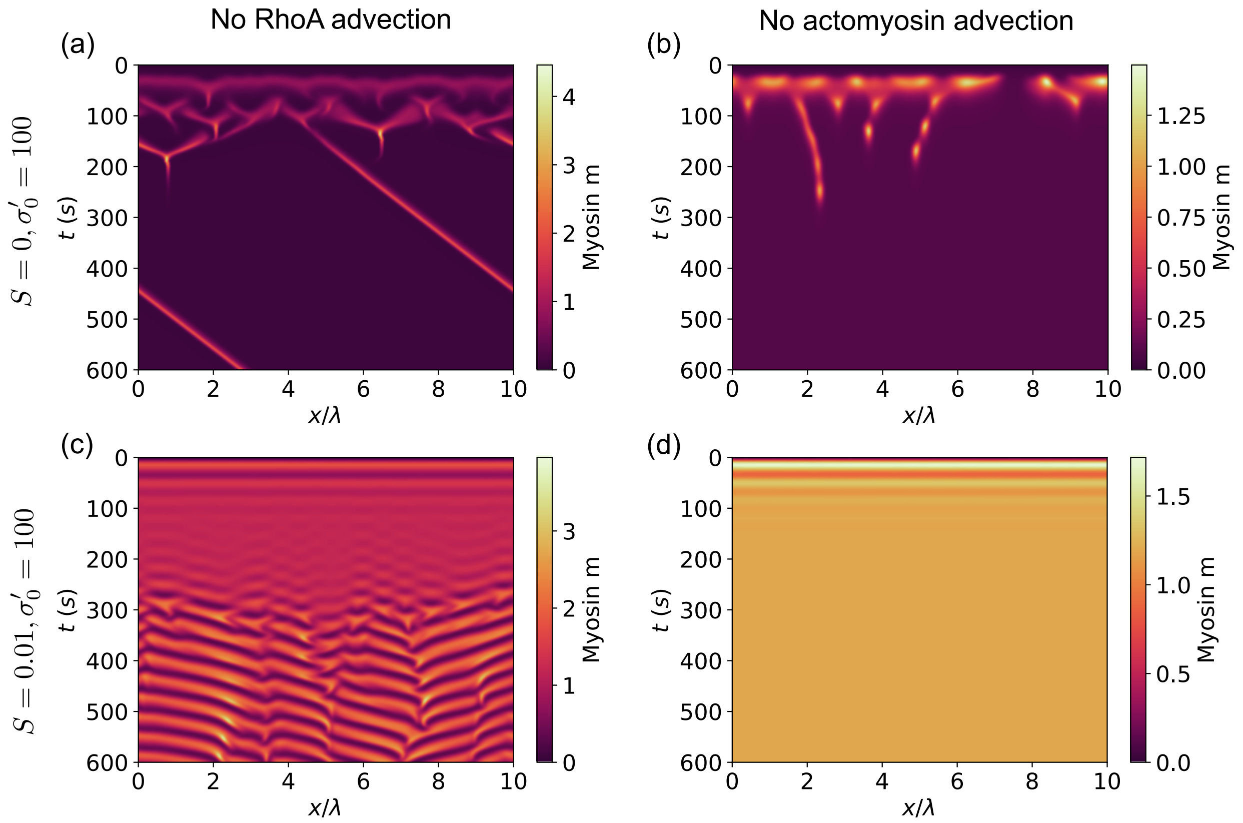


**Figure S 4. RhoA and actomyosin advection are necessary for stable pattern formation and propagating waves.** (a-d) Kymographs of actomyosin concentration with no (a, c) RhoA advection , and (b, d) no actomyosin advection , for (a, b) $S = 0.00\ \mathrm{s}^{-1}$, $\sigma'_a = 100$, and (c, d ) $S = 0.01\ \mathrm{s}^{-1}$, $\sigma'_a = 100$.

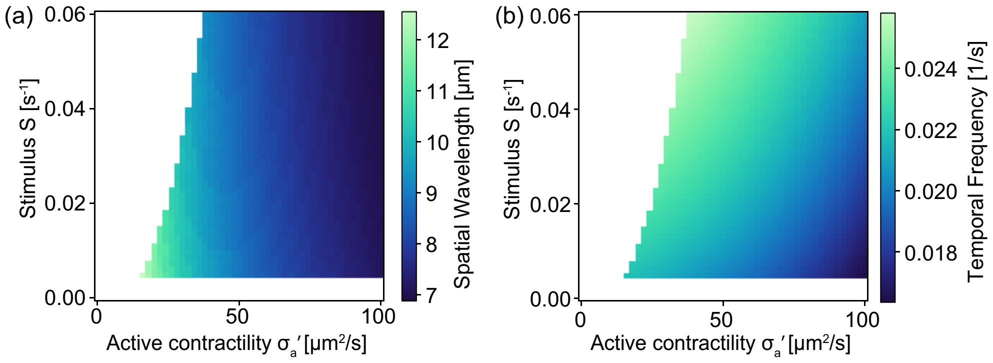


**Figure S** 5. **Linear stability analysis predicts the frequency and wavelength of pulsatile contractins.** (a) Spatial wavelength, and (b) temporal frequency computed by linear stability analysis for varying active contractility $\sigma_a'$ and RhoA stimulus $S$.
